# Interdependence of balance mechanisms during bipedal locomotion

**DOI:** 10.1101/658120

**Authors:** Tyler Fettrow, Hendrik Reimann, David Grenet, Elizabeth Thompson, Jeremy Crenshaw, Jill Higginson, John Jeka

**Affiliations:** Kinesiology and Applied Physiology, University of Delaware, Newark, DE 19713, United States of America; Department of Physical Therapy, University of Delaware, Newark, DE 19713, United States of America; Department of Kinesiology, Temple University, Philadelphia, PA 19122, United States of America; Department of Physical Therapy, Temple University, Philadelphia, PA 19122, United States of America; Department of Mechanical Engineering, University of Delaware, Newark, DE 19713, United States of America

## Abstract

Our main interest is to identify how humans maintain upright while walking. Balance during standing and walking is different, primarily due to a gait cycle which the nervous system must contend with a variety of body configurations and frequent perturbations (i.e., heel-strike). We have identified three mechanisms that healthy young adults use to respond to a visually perceived fall to the side. The lateral ankle mechanism and the foot placement mechanism are used to shift the center of pressure in the direction of the perceived fall, and the center of mass away from the perceived fall. The push-off mechanism, a systematic change in ankle plantarflexion angle in the trailing leg, results in fine adjustments to medial-lateral balance near the end of double stance. The focus here is to understand how the three basic balance mechanisms are coordinated to produce an overall balance response. The results indicate that lateral ankle and foot placement mechanisms are inversely related. Larger lateral ankle responses lead to smaller foot placement changes. Correlations involving the push-off mechanism, while significant, were weak. However, the consistency of the correlations across stimulus conditions suggest the push-off mechanism has the role of small adjustments to medial-lateral movement near the end of the balance response. This verifies that a fundamental feature of human bipedal gait is a highly flexible balance system that recruits and coordinates multiple mechanisms to maintain upright balance during walking to accommodate extreme changes in body configuration and frequent perturbations.

## 1. Introduction

Investigations of human balance control have focused primarily on standing, either during unperturbed quiet stance (Winter et al., 1998; Wang and Newell, 2014) or in response to mechanical (Horak and Nashner, 1986) and sensory perturbations (Peterka, 2002; Oie et al., 2002). Surprisingly, the literature on balance control during walking is relatively sparse, despite the fact that most falls occur during walking (Robinovitch et al., 2013; Tinetti et al., 1988; Berg et al., 1997). One explanation for this disparity is that walking entails many behaviors that are not relevant to standing, fostering investigations into navigation (Patla and Greig, 2006; Jansen et al., 2011), obstacle avoidance (Rietdyk and Rhea, 2011) and metabolic efficiency (Maxwell Donelan et al., 2001). However, we argue that balance control during walking requires an analysis that differs markedly from standing balance, for a number of reasons. First, the temporal window over which the human body propels itself (i.e., the gait cycle) allows for distinct balance responses to be initiated sequentially. Second, body configuration changes dramatically over the gait cycle (e.g., single stance to double stance), resulting in balance responses that are highly constrained by this reconfiguration. Third, the nervous system must respond not only to external disturbances while walking but respond to disturbances that are inherent to the gait cycle itself (e.g., heel-strike). In this regard, walking is often conceived as a hybrid dynamical system, combining elements of both continuous and discrete dynamics, requiring a different form of control than standing. Here we determine how the human nervous system coordinates multiple balance mechanisms during walking to determine whether such mechanisms are inter-dependent as they unfold over the gait cycle.

We have identified three basic mechanisms following visual perturbations in the medial-lateral direction that shift the CoP in the desired direction, the lateral ankle mechanism, the foot placement mechanism, and the push-off mechanism (Reimann et al., 2018b). We provide a description of these basic control mechanisms available to healthy adults, and determine whether the mechanisms are temporally coordinated throughout the gait cycle. We expect all three mechanisms to be interdependent to produce an overall balance response.

The lateral ankle mechanism refers to the generation of ankle inversion/eversion torque during the single stance phase. Rolling the ankle while the foot is on the ground can shift the CoP under the foot. The goal is to shift CoP in the direction of the perceived fall under the stance foot. The lateral ankle mechanism is able to act on the CoP at virtually any time during the gait cycle, and is the main means of modulating the CoP under the stance foot during sustained locomotion. However, the magnitude of CoP modulation that the lateral ankle mechanism can produce is constrained to the width of the foot or shoe.

The foot placement mechanism is another mechanism that can shift the CoP. The swing foot is shifted in the direction of the perceived fall, and on heel-strike, shifts the CoP. Foot placement is the most commonly reported measure of dynamic stability (Young and Dingwell, 2013; Yiou et al., 2017). Much of the balance and walking literature focuses on swing foot placement in terms of overall step width or step width variability (Schrager et al., 2009; Brach et al., 2005; Hurt et al., 2010). The foot placement mechanism is different from step width, which assumes an increase in step width. Here we are interested in the control of foot placement in order to modulate the CoP in a particular direction, which can result in an increased or a decreased step width.

The push-off mechanism refers to the modulation of ankle plantarflexion angle during double stance. Very few studies have recognized the push-off mechanism’s role in balance. Ankle plantarflexion torque has been shown to change as a result of bipolar, binaural galvanic vestibular stimulation (Iles et al., 2007) and anterior-posterior mechanical perturbations (Vlutters et al., 2016). Only recently has the push-off been verified to have a functional role in balance control in the medial-lateral direction (Kim and Collins, 2015; Reimann et al., 2018a). In response to a visually perceived fall to the side, we observed a direction-dependent modulation of the stance leg ankle plantar/dorsiflexion angle (Reimann et al., 2018b).

We believe these three mechanisms are the main available methods to make fine motor adjustments to the CoP and CoM in order to maintain balance while walking. We believe this because these were three separate biomechanical responses to the visually perceived fall that are functionally separate from one another and neurally driven (Reimann et al., 2018b). Given that multiple mechanisms are available to the CNS to maintain balance, we analyzed whether these mechanisms are functionally related. For example, modeling results suggest that without any CoP modulation during the stance phase following a sensory perturbation, a 60% larger foot placement is needed to maintain balance in the medial-lateral direction (Reimann et al., 2017). Here we determine if the human nervous system coordinates the balance response throughout the gait cycle by testing whether these mechanisms are interdependent to produce an overall balance response.

## 2. Methods

Portions of this data and detailed data collection methods have already been published (Reimann et al., 2018b), where we focused entirely on group averages. Here we focus on the balance mechanisms on every step. Twenty healthy young subjects (11 female, 22.8 ± 4.1years, 75.2 ± 17.9kg) volunteered for the study. Subjects provided informed written consent to participate. Subjects did not have a history of neurological disorders or surgical procedures involving legs, spine or head. The experiment was approved by the Temple University Institutional Review Board.

### 2.1. Experimental design

After explaining the experiment, obtaining consent and placing markers and EMG sensors, subjects first walked for 15 min on the self-paced treadmill in the virtual environment to adapt to this experimental setup. We then stopped the treadmill and told the subjects that we would now perturb their sense of balance by modifying the virtual scene, and asked them to cope with this perturbation normally and keep walking forward. Data collection blocks consisted of two alternating phases for metronome and stimulus. During metronome phases, lasting 30 s, subjects were provided an auditory metronome at 90 bpm and asked to use this as an approximate guideline for their footsteps, both during metronome and stimulus phases. During stimulus phases, lasting 120 s, the metronome was turned off, and subjects received visual fall stimuli as described above. Data were collected during stimulus phases. Each subject performed four blocks of walking, each block consisting of five metronome and five perturbed phases, always starting with metronome phases, for a total of 12.5 min per block. After each block, the treadmill was turned off and subjects were offered a break. This protocol was implemented in a custom Labview program that sent the head position, treadmill speed and rotation angle to the Unity computer via UDP and saved the visual rotation angle and treadmill speed at 100Hz.

Subjects walked on a split-belt, instrumented treadmill within a virtual environment projected onto a dome surrounding the treadmill by ∼180 degrees (Bertec, Inc.). The treadmill was self-paced, using a nonlinear PD-controller in Labview (National instruments Inc., Austin, TX, USA) to keep the MPSIS markers on the mid-line of the treadmill. Perturbations consisted of a rotating virtual environment scene at a rate of 60° s^−2^ for 600 ms around the anterior-posterior axis of the midline of the floor, inducing a feeling of falling to the side. The perturbations were provided randomly on heel-strike every 10-13 steps (stimulus trigger), and rotation direction randomized to produce the perception of falling to the left or right. The resulting rotation of 10.8° was then held constant for 2,000 ms, before being reset to neutral rotation with uniform speed over 1,000 ms. After resetting to neutral rotation, a randomized interval of 1013 steps elapsed before the next stimulus was triggered. Heel strikes were identified as downward threshold crossings of the vertical heel-marker position. The threshold was set to the vertical heel-marker position of each foot during quiet standing, plus 3mm.

Reflective markers were placed bilaterally on the feet, lower legs, thighs, pelvis, torso, head, upper arms, forearms, and hands of the subject, using the Plug-in Gait marker set (Davis et al., 1991) with six additional markers on the anterior thigh, anterior tibia, and 5th metatarsal of each foot for a total of 45 markers. Marker positions were recorded at 250Hz using a Vicon motion capture system with nine cameras. Ground reaction forces and moments were collected at 1,000 Hz from both sides of the instrumented split-belt treadmill (Bertec, Inc.). Forces and moments were transformed into a common coordinate frame and then used to calculate the whole-body CoP (Winter et al., 1990).

### 2.2. Data Management and Organization

Kinematic data were low pass filtered with a 4th order Butterworth filter at a cut-off frequency of 10Hz. Small gaps in the marker data of up to 100ms length from occlusions were filled using cubic splines. Time points with remaining marker occlusions were excluded from further analysis. From the marker data, we calculated joint angle data based on a geometric model with 15 segments (pelvis, torso, head, thighs, lower legs, feet, upper arms, forearms, hands) and 38 degrees of freedom (DoF). We estimated the hip joint centers based pelvis landmarks (Tylkowski et al., 1982; Bell et al., 1990), and the knee joint centers and knee flexion rotational axes from reference movements using the symmetrical axis of rotation approach (Ehrig et al., 2007). We performed inverse kinematics by minimizing the distance between the measured and the model-determined marker positions (Lu and O’Connor, 1999). This optimization was performed first for the six pelvis DoFs, which formed the root of the kinematic tree, then for the six DoFs at the lumbar and cervical joints, and last for each of the arms and legs separately. We estimated the body center of mass (CoM) positions based on estimated segment CoM locations (Dumas et al., 2007) and the inverse kinematics and calculated CoM velocities and accelerations using numerical derivation by time.

We identified heel strike events for each foot by finding negative peaks in the vertical positions of the heel markers with minimal inter-peak distances of 250 ms and peak prominence greater than 2 cm, and push-off events as the first peak in the vertical velocity of the 2nd metatarsal marker with a prominence greater than 0.35 ms^−1^ after each heel strike. We visually inspected the result of this automatic identification and applied manual corrections in the rare cases where events were misidentified.

The experimental design had 4 distinct stimulus conditions, where each leg could trigger a stimulus to the left or right. All data between heel-strikes were normalized to 100 time steps. We subtract the *control* mean from all data, including *control*, for every subject. The control steps are defined as the two steps preceding the heel-strike that triggers the visual perturbation. All data between heel-strikes were normalized to 100 time steps. All of the data here are represented as a change in response from the average of the *control* steps.

#### Lateral Ankle Mechanism

The lateral ankle mechanism is defined as the integrated response from *control* steps of the medial-lateral distance between CoP and CoM during single stance. The integration is performed during single stance to avoid the contribution of the foot placement mechanism on the CoP. Throughout the text we refer to this variable as Ankle.

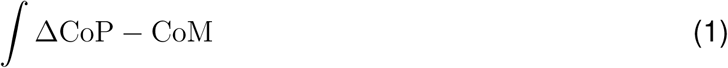

#### Foot Placement Mechanism

We define the foot placement mechanism by the medial-lateral response of the swing leg heel position at heel strike, relative to the initial medial-lateral position of the trigger foot heel strike. Throughout the text we refer to this variable as Foot.

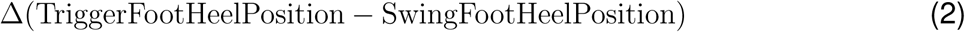

#### Push-off Mechanism

We define the push-off mechanism as the integrated response from *control* steps of the trigger foot ankle dorsiflexion/plantarflexion angle during the second double stance phase following stimulus trigger. Throughout the text we refer to this variable as Push.

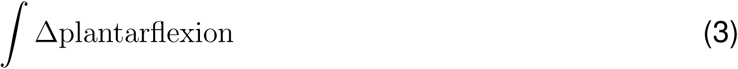

The experimental design had four distinct stimulus conditions, where each heel strike could trigger a stimulus to the left or right. Here we assume anatomical symmetry and group the conditions that are anatomically similar (i.e. [Right heel strike triggers Stimulus Right, Left heel strike triggers Stimulus Left]). Previous analysis of the data set indicate asymmetries between feet are not present (Reimann et al., 2018b). We also analyzed how the mechanisms are coordinated during *control* steps. This leaves three conditions to analyze, stimulus *towards* trigger foot, stimulus *away* from trigger foot, and *control*. After processing, we were left with 1947 steps for *Stimulus Towards*, 1930 steps for *Stimulus Away*, and 3854 steps for *Control*.

### 2.3. Statistical Analysis

We confirmed the assumptions of normality and homoscedasticity by visual inspection of the residual plots for each combination of variables. Linear mixed models were used for each pair of mechanisms to assess interdependence. For each pair of mechanisms, we fitted a linear mixed model and performed an ANOVA to analyze the interdependence of the mechanisms and interaction of stimulus direction, using Satterthwaites method (Fai and Cornelius, 1996) implemented in the R-package lmerTest (Kuznetsova, 2017). We used R (R Core Team, 2013) and lme4 (Bates et al., 2009) to assess the significance of slopes of the regressions for each pair of mechanisms. Confidence intervals were calculated for the slopes for each stimulus direction using the function confint. The outcome and predictor variables for each model were chosen based on temporal order (i.e. Ankle is used prior to Foot, and Foot prior to Push). We included stimulus direction as a predictor and allowed random intercepts by Subject and random slopes. The following model is an example for assessment of each pair of mechanisms:

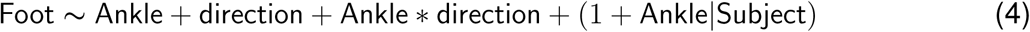

For visual purposes, we perform a least-squares fit for all pairs of mechanisms. Pearson Correlation R^2^ values were calculated for each balance mechanism pair to provide an indication of the strength of the relationship.

## 3. Results

All figures show data averaged from eighteen out of twenty subjects. We removed two subjects from this analysis. These two subjects expressed R^2^ values two standard deviations lower than the mean for Ankle-Foot interdependence. Including the subjects resulted in non-converging of the mixed-models in R.

The balance response to a perceived fall laterally (towards and away from trigger foot) consists of a combined response of the lateral ankle mechanism, the foot placement mechanism and the push-off mechanism, as shown in Figure 1. All mechanisms are shown as time series to illustrate their temporal order. After onset of the stimulus on the trigger foot heel strike, the lateral ankle mechanism is the first to act (Figure 1A), on average approximately 350 ms post-stimulus where the *Stimulus Towards* and *Stimulus Away* deviate away from each other. The activation of the lateral ankle musculature leads to a CoP shift towards the direction of the perceived fall, for both *Stimulus Towards* and *Stimulus Away*. Figure 1B shows the initiation of the foot placement mechanism begins around 450 ms, represented by the change in swing foot heel position deviating from the control steps towards the end of single stance. This shift starts prior to heel strike, but does not substantially affect the CoP until heel strike. The foot is placed in the direction of the perceived fall. The push-off modulation begins around mid-stance (∼450-550 ms), but the majority of the angle change occurs during the second double stance following the stimulus onset (Figure 1C). We observed a plantarflexion response for *Stimulus Towards* and a dorsiflexion response for *Stimulus Away*.

**Figure 1:**
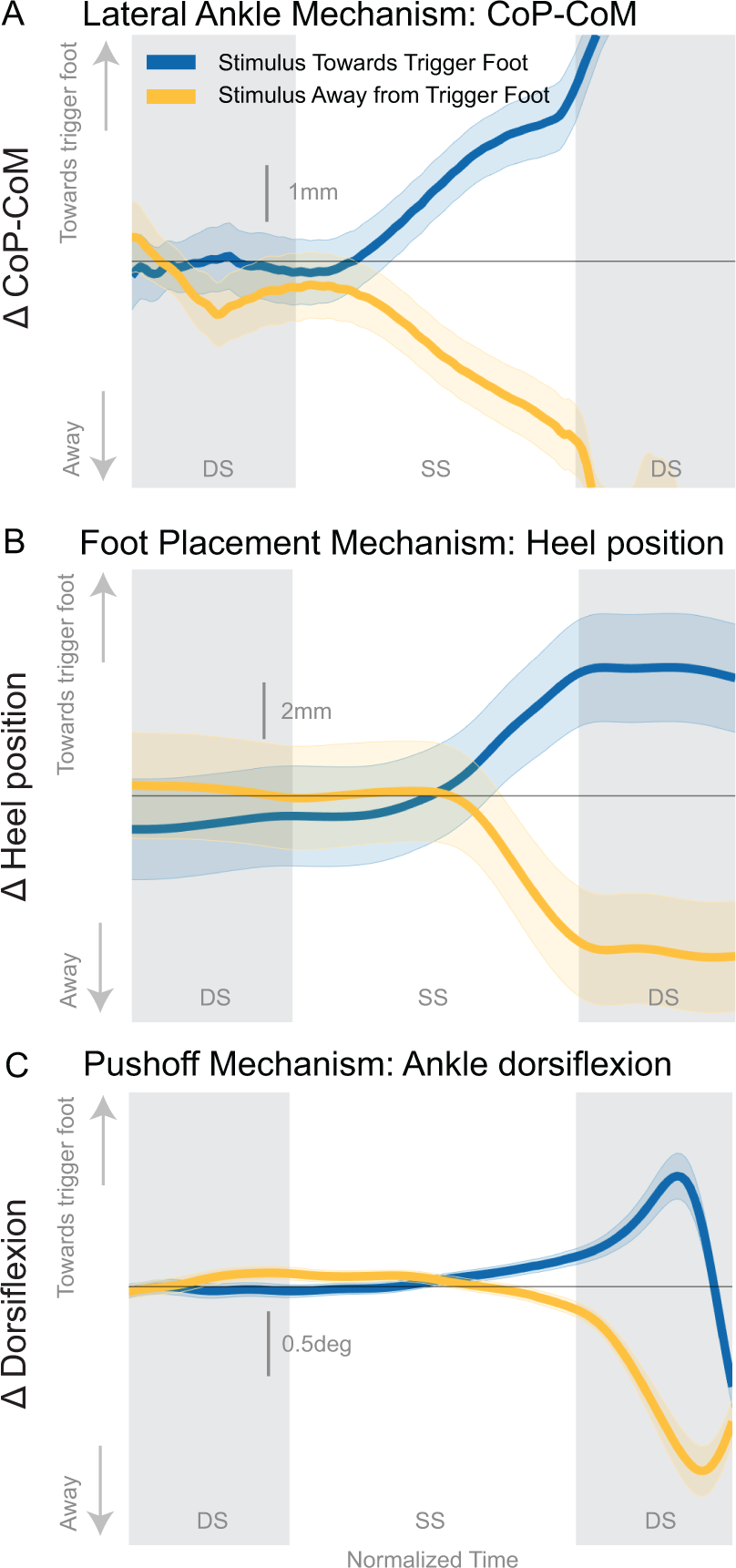
The average difference from control (Δ or response) for three major balance mechanisms are displayed for Stimulus Towards (Blue) and Stimulus Away (Yellow) encased by the 95% confidence interval. Double stance (DS) is displayed as the grey shaded areas.

### 3.1. Interdependence

Figure 2 illustrates the interdependence between pairs of mechanisms by displaying the use of each mechanism for a given gait cycle as individual data points. The correlation between the foot placement and the lateral ankle mechanism is the strongest relative to the other mechanism pairs (Figure 2A) for *Stimulus Towards, Stimulus Away* and *Control*, demonstrating a negative interdependent relationship for all conditions (see Table 1, 2). A larger response of the lateral ankle results in a smaller foot placement response (and vice versa). A positive but weak correlation exists between the push-off and foot placement mechanisms, for *Stimulus Towards, Stimulus Away*, and *Control*, Figure 2B), suggesting that Foot Placement and Push-off are also interdependent (see Table 1, 2). Smaller foot placement leads to smaller push-off (and vice versa). Finally, the push-off and lateral ankle mechanisms demonstrate a negative trend *Stimulus Away* (see Figure 2C and Table 2). *Stimulus Towards* and *Control* did not have slopes significantly different from zero, yet trended towards a negative relationship, suggesting the lateral ankle and push-off mechanisms are very weakly interdependent.

**Table 1:** Results of the Mixed Model output indicating which factors have a significant effect on the outcome variable or dependent variable (see Statistic Analysis section 2.3.)

|  | Fixed-Effect | Numerator Df | Denominator Df | F | p |
| --- | --- | --- | --- | --- | --- |
| Foot | Ankle | 1 | 18 | 792.47 | <0.0001 |
|  | Direction | 2 | 7710 | 178.57 | <0.0001 |
|  | Ankle*Direction | 2 | 7684 | 2.98 | 0.051 |
| Push | Ankle | 1 | 17 | 7.59 | 0.0133 |
|  | Direction | 2 | 7711 | 138.27 | <0.0001 |
|  | Ankle*Direction | 2 | 7714 | 0.36 | 0.697 |
| Push | Foot | 1 | 17 | 34.15 | <0.0001 |
|  | Direction | 2 | 7708 | 106.27 | <0.0001 |
|  | Foot*Direction | 2 | 7712 | 1.77 | 0.171 |

**Table 2:** Estimated 95% confidence interval of the slope of regression are displayed in the Lower and Upper columns. Confidence intervals that exclude zero are made bold and deemed a significant relationship.

| Regression Tested | Stimulus Towards |  | Stimulus Away |  | Control |  |
| --- | --- | --- | --- | --- | --- | --- |
|  | lower | upper | lower | upper | lower | upper |
| Ankle(mm s)-Foot(mm) | **-6.94** | **-5.34** | **-6.2** | **-5.27** | **-6.762** | **-5.261** |
| Foot(mm)-Push(deg s) | **0.0018** | **0.007** | **0.003** | **0.006** | **0.001** | **0.006** |
| Ankle(mm s)-Push(deg s) | -0.029 | 0.006 | **-0.024** | **-0.003** | -0.031 | 0.002 |

**Figure 2:**
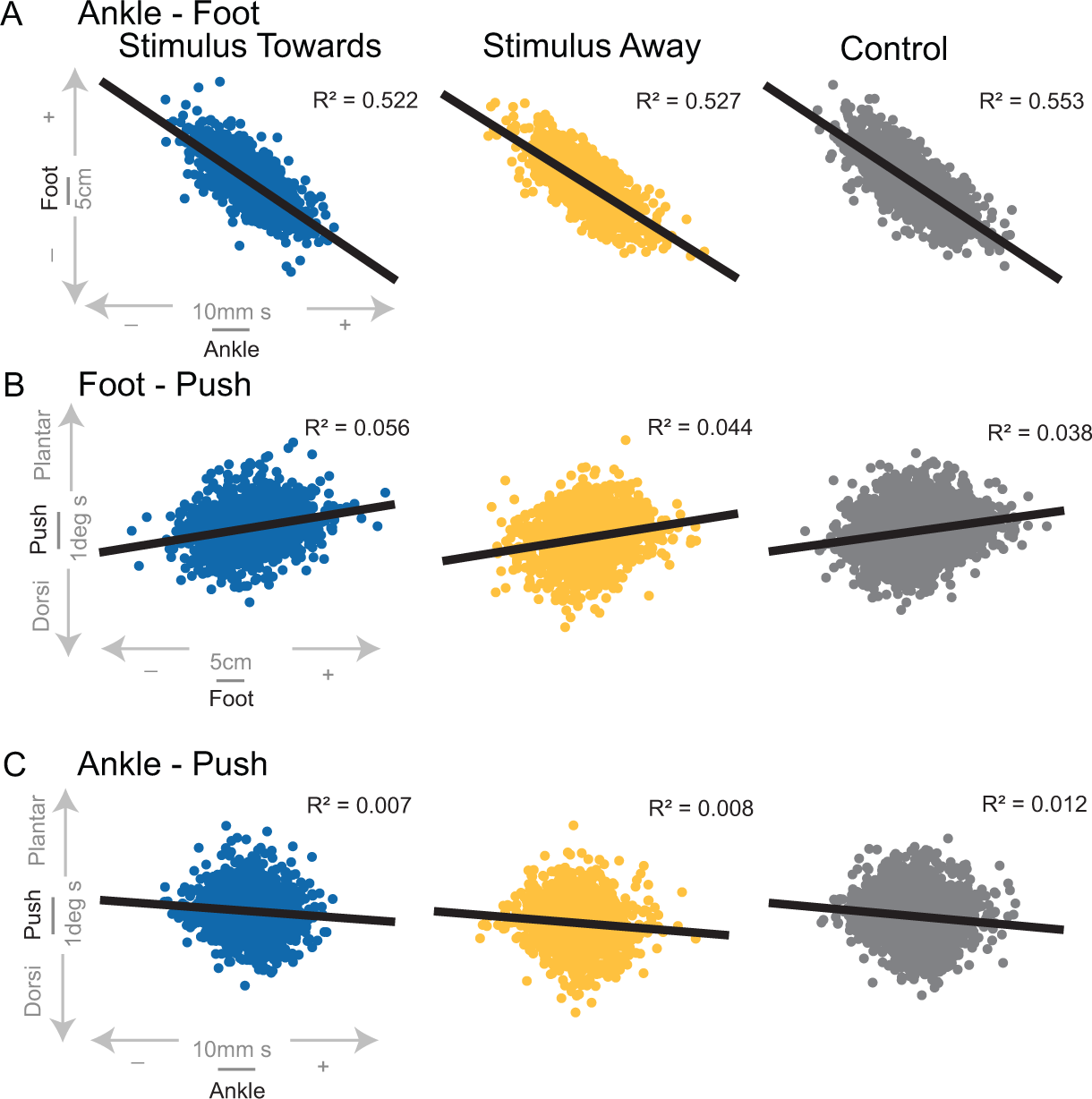
The relationship between the three major balance mechanisms are displayed for each stimulus condition with least-squares fit and corresponding Pearson correlation R^2^ values. Each data point represents the combination of the balance mechanisms displayed on the axes for that figure. Values towards + imply a response in the direction towards the trigger foot.

The point clouds of the balance mechanisms across the three stimulus directions in Figure 2 are difficult to distinguish visually. When we remove individual data points from Figure 2 and only include the means and least-squares fits in Figure 3, a clear separation of the stimulus conditions is evident. Balance responses are toward the trigger foot, relative to control steps, when the stimulus is towards the trigger foot. Likewise, in the *Stimulus Away* conditions, balance responses are further away from the trigger foot compared to control steps. This effect of stimulus direction (confirmed by significance of fixed-effect term Direction in first three mixed-models (Table 1), consistent with results previously published by Reimann et al. (2018b). The fixed-effect term Ankle*Direction tests the change in slope of the models by stimulus direction. No mechanism pair had significantly different slopes across condition, including Ankle-Foot, Ankle-Push, and Foot-Push (see Table 1).

**Figure 3:**
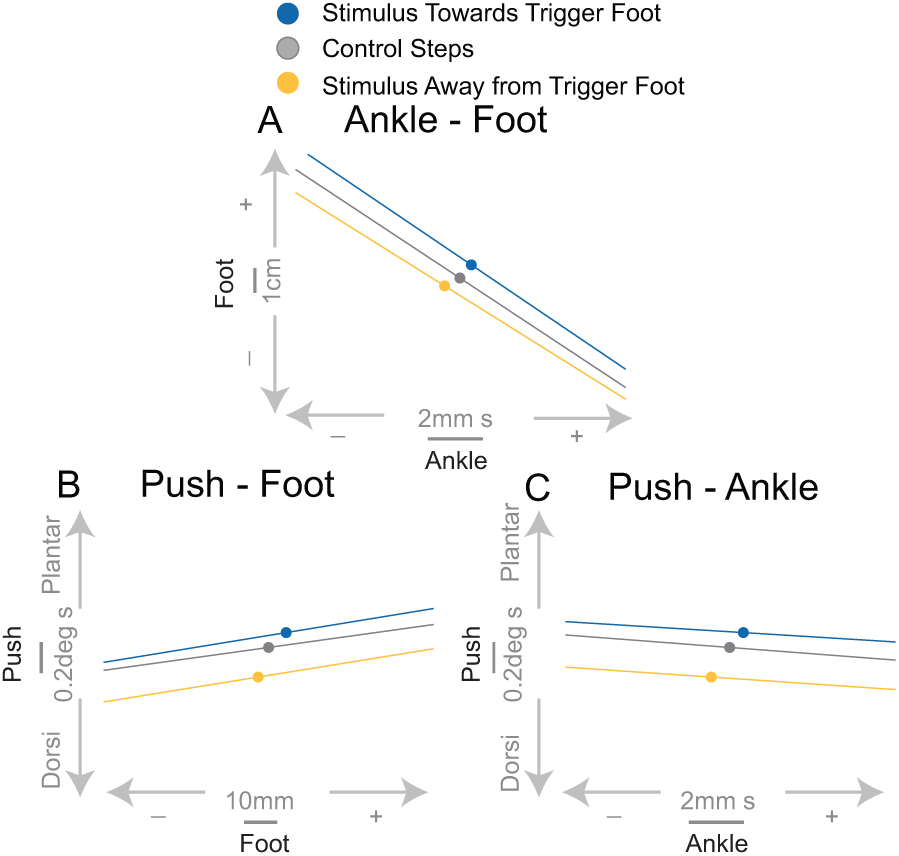
The average for each combination of balance mechanisms for each condition are displayed as solid dots. The least-squares fits are also displayed. Values towards + imply a response in the direction towards the trigger foot.

## 4. Discussion

There is a high degree of variability in human gait, even in the control of balance, but we show that the nervous system is able to overcome this variability by making adjustments throughout the gait cycle through the coordination of balance mechanisms. Considering the large number of degrees of freedom involved in bipedal locomotion and resultant body configuration that varies with each step, flexible implementation of balance mechanisms may be fundamental to maintain upright during walking.

We have identified three basic balance mechanisms that are used to respond to a perceived fall to the side during locomotion as well as during unperturbed walking. The average balance response as shown in Figure 1 suggests the use of all mechanisms regardless of stimulus direction. The lateral ankle mechanism modulates the center of pressure (CoP) under the stance foot. Change in foot placement then moves the CoP after a modulation of foot position during the swing phase of the gait cycle. We also observed a change in ankle plantarflexion angle depending on the direction of the stimulus within the first gait cycle, indicating that push-off modulation is another mechanism used to respond to a perceived fall to the side. Our conclusions are: 1) balance mechanisms during walking are interdependent; 2) the push-off mechanism is used to make subtle adjustments to balance, but is not strongly interdependent with the other mechanisms; and 3) the use of each balance mechanism alone may be inaccurate (as evidence by common overshooting and undershooting of the lateral ankle mechanism), but the nervous system is able to adjust by using a variety of balance mechanisms throughout the gait cycle.

### 4.1. Interdependence

The combination of the mechanisms throughout the gait cycle have a compounding effect on balance. Balance throughout the gait cycle may be conceived as a zero-sum scenario. If a corrective action is performed early in the gait cycle, it reduces the need for a corrective action later in the cycle. The focus of this study was to unpack how all three mechanisms are interdependent.

All pairs of balance mechanisms are significantly correlated with one another, despite small R^2^ values, according to the linear mixed model results. Two relationships were negatively correlated: lateral ankle - foot placement and push-off - foot placement. With these two negative relationships, it follows that foot placement and push-off are positively related. Given the disparity in the strength of the correlations between balance response pairs, it seems that the initial response to a disturbance is critical to the overall balance response, leaving the push-off to play a small but systematic modulating role that does not depend strongly on the responses previous to its initiation. Although an interaction between the lateral ankle and foot placement mechanism has been observed previously (Hof et al., 2007, 2010), these reports suggest the lateral ankle mechanism adjusts inaccuracies in foot placement. The current results suggest that foot placement also adjusts to inaccuracies in the lateral ankle mechanism, highlighting an interdependent relationship. The lateral ankle mechanism commonly overshoots or under-shoots the required modulation, as evidence by the range of values observed in Figure 2A. This could be viewed as inaccurate use of the lateral ankle mechanism, or be a result of the natural variability in gait and adjustment to the preceding inaccurate foot placement. The subsequent mechanisms (foot placement and push-off) are able to adjust for this inaccuracy or variability.

The current results indicate that there is a third mechanism involved in the coordination between balance mechanisms. The push-off is typically thought to contribute to propulsion, but we believe the push-off is also used as a subtle adjustment to medial-lateral balance, for two reasons. 1) We observe a systematic modulation of ankle plantarflexion angle following a medial-lateral (ML) balance perturbation. It seems unlikely that a response to an ML perturbation would serve only propulsion in the sagittal plane. 2) Propulsion is dependent on the stimulus direction (Figure 1C). If, for example, propulsion serves to improve stability, we would expect an increase in propulsion regardless of the stimulus direction. However, only when the perturbation is towards the trigger foot, is there an increased push-off. We see a decreased push-off (i.e. plantarflexion) when the perturbation is away from the trigger foot, suggesting a decrease in propulsion that results in an adjustment of the CoM over the base of support. Furthermore, Klemetti et al. (2014) has provided evidence that ankle planterflexion torque induces trunk roll accelerations, providing further support for the push-off acting in the medial-lateral direction. Only recently has the push-off been verified to have a functional role in balance control in the medial-lateral direction as Kim and Collins (2015) show modulation of ankle torque based on CoM behavior can reduce metabolic expenditure. We speculate that the push-off is mostly acting in the anterior-posterior direction, but the offset of the feet laterally during double stance could allow for a rotation about the moment arm between the trailing limb and the CoM. Thus, the subtle modulation of push-off is less about moving the body forward and more about shifting weight to the alternate foot.

Despite the systematic nature of the push-off, the R^2^ values between push-off and other mechanisms were very small. We can provide three possible reasons for such small but consistent relationships: 1) The push-off mechanism may provide more of an indirect balance adjustment through modulation of other gait parameters, which we are currently analyzing and plan to address in a future publication; 2) The relationship between the lateral ankle and foot placement mechanisms is so strong that it masks their relationship with the push-off mechanism; 3) There are other mechanisms contributing to the maintenance of balance during walking. As discussed in the previous paragraph, we are confident the push-off is a separate balance mechanism, despite the weak correlations, in that it is activated separately from the lateral ankle and foot placement mechanisms. The small correlations between push-off and the other two balance mechanisms is actually evidence that they are separately activated balance mechanisms. For example, if the lateral ankle and push-off were activated through the same neural pathway, we would expect a strong positive relationship between the two. We actually observe a weak negative correlation between the lateral ankle and push-off mechanisms. The weak correlations may also be a product of the flexibility of the healthy young nervous system. There are so many degrees of freedom available to act on the balance related forces that a moderate amount of error at any point of the gait cycle is acceptable.

An interesting finding was that these balance responses are observed on steps both with and without perturbations. This is consistent with the view that heel strike is essentially a perturbation that is inherent to the gait cycle, requiring a control action to maintain upright balance on every step (Bauby and Kuo, 2000; Wang and Srinivasan, 2014). This finding is substantiated by the fact that the slopes of each balance mechanism pair are not statistically different across all three stimulus directions. (*Stimulus Towards, Stimulus Away, Control)*. Thus, interdependence of the balance mechanisms is present in all conditions, but the overall balance response is shifted in the direction of the perceived fall. This emphasizes that a sensory perturbation does not alter the fundamental nature of the balance response observed during unperturbed walking, but in the presence of a perceived threat to balance, the overall balance response shifts in the direction of the perceived fall (i.e. blue shifts towards the trigger foot and yellow shifts away from trigger foot in Figure 3).

### 4.2. Limitations

Previous work has identified different strategies that can be recruited to aid in the overall goal of maintaining stability. Otten (1999) has shown through simulations of narrow beam walking that rotations of the arms, neck, and hips can limit the angular accelerations of the head-arms-trunk complex, and thus the CoM. Rebula et al. (2017) reports that turning (i.e. foot yaw rotation) can have the same effect on medial-lateral balance as a lateral foot placement. A foot yaw response may also have an impact on the function of a push-off response, as plantarflexion through different ranges of an externally rotated foot will create different directional ground reaction forces. We did not observe such responses in the current experiment, suggesting that there may be particular situations that lead to less common but effective balance responses.

We also assume all of the modulation of the CoP with respect to the CoM in the first single stance following the trigger perturbation is a result of the lateral ankle mechanism, but it is certainly possible other mechanisms could be used to assist in this modulation. Hof and Duysens (2018) also found that most of the CoP modulation is a result of ankle muscle activation, but acknowledge that the ankle responses to a perceived fall laterally is not a simple inversion/eversion activation. Future work will seek to uncover other mechanisms that can contribute to the modulation of the CoP with respect to the CoM.

We note a change in ankle plantarflexion in response to the medial-lateral balance perturbation; however, we do not have concrete evidence that the change in ankle plantarflexion angle is due to an increased ankle plantarflexion torque and propulsive ground reaction force. Due to the inherent design of our experiment, calculation of joint torques and ground reaction forces from a single limb are impossible to discern because of scenarios involving one foot on two force plates or two feet on one force plate. We asked people to treat the split-belt treadmill as a sidewalk to create an environment the allows for the most natural gait. Adding medial-lateral balance perturbations increases the number of instances where a single foot may not land squarely on a single force plate. This scenario makes it difficult to provide concrete evidence that the ankle plantarflexion response is producing a “push-off”. Future work will investigate in detail how the systematic change in ankle plantarflexion angle produces medial-lateral balance shifts, but for now we believe there is enough support to assume it does produce a CoM shift in the medial-lateral direction.

### 4.3. Conclusion

In conclusion, an interdependent relationship has been uncovered between three basic mechanisms for the maintenance of upright balance during walking. Most evidently, our analysis illustrates that when the lateral ankle mechanism overshoots or undershoots the foot placement will provide a correction later in the gait cycle. The push-off mechanism is not as strongly related to the other two balance mechanisms described, suggesting the push-off is only producing a subtle balance related adjustment. Moreover, we highlight that these mechanisms are used on every step, regardless of whether there is a perturbation. The perturbations result in a shift of the overall balance response in the direction of the perceived fall. This is confirmation that a fundamental feature of human bipedal gait is a highly flexible balance system that recruits multiple mechanisms to maintain upright balance during walking despite extreme changes in body configuration and frequent perturbations.

